# Fluid replacement and acute systemic inflammatory response syndrome during prolonged eight-hour extreme heat exposure in young men

**DOI:** 10.1101/2024.12.12.628082

**Authors:** Faming Wang, Caiping Lu, Tze-Huan Lei, Ying Lei

## Abstract

Prolonged exposure to extreme heat poses significant risks, including systemic inflammatory response syndrome (SIRS), organ damage and hormonal imbalance. While fluid replacement is commonly recommended to mitigate these effects, its efficacy under uncompensable heat stress remains unclear. This study investigated the impacts of fluid replacement on thermoregulation, systemic inflammation, organ stress, cortisol levels and plasma electrolyte balance during eight-hour of extreme heat exposure in healthy young men. Twelve participants (age: 24.7±1.6 years; body surface area: 1.9±0.1 m²) underwent two randomized trials (dehydration: 125 mL/hour; euhydration: 375 mL/hour) in a heat chamber (40 °C, 55% RH). Biomarkers of inflammation (e.g., IL-6, IL-1β), oxidative stress (e.g., MDA, SOD), organ function (ALT, BUN), cortisol, and electrolytes (sodium, potassium, chloride) were measured before and after exposure. Core temperature (*T_core_*) was continuously monitored. Results showed that fluid replacement significantly reduced *T_core_* at the end of the exposure (38.0±0.12 °C vs. 38.2±0.10 °C, *p*=0.046). However, it exacerbated systemic inflammation (IL-6: euhydration 19.8±4.3 pg/mL vs. dehydration 12.5±2.8 pg/mL, *p*<0.01) and liver stress (ALT: euhydration 45.3±6.7 U/L vs. dehydration 34.1±5.5 U/L, *p*=0.03). Cortisol levels decreased significantly in the euhydration group (*p*=0.041), potentially indicating attenuated stress resilience. Electrolyte imbalances (reduced sodium and potassium concentrations) were observed in the euhydration condition. Taken together, while fluid replacement reduced *T_core_*, it did not mitigate SIRS and instead exacerbated systemic inflammation, liver stress, and electrolyte imbalances, potentially through hypotonic osmotic stress. These findings underscore the need for personalized hydration strategies that balance fluid and electrolyte intake during extreme heat exposure to minimize health risks.

**Key points:**

- This study investigated the effects of fluid replacement on acute inflammatory responses, thermoregulation, and organ stress markers in healthy young males exposed to eight hours of uncompensable heat stress.
- Fluid replacement did not mitigate acute systemic inflammatory response syndrome (SIRS) but exacerbated inflammatory and organ stress markers despite reducing core temperature.
- Core temperatures remained significantly lower in the euhydration group, but systemic and hepatic inflammatory markers worsened, highlighting the complexity of hydration strategies under extreme heat.
- We emphasize the need for individualized hydration strategies incorporating electrolyte balance and sweat rate monitoring to minimize SIRS risk.
- We provide actionable insights into heat stress physiology, with implications for occupational health guidelines, public health policies, and climate change adaptation.

## Introduction

Heatwaves have become more intense and frequent in recent years, impacting countries worldwide (Meehl & Tebaldi, 2004). Animal studies have shown that extreme heat exposure increases core temperature (*T_core_*) and places considerable cellular stress on tissues (Welc *et al*. 2013), which in turn activates heat shock factor 1 (HSF1) gene transcription (Winter-Vann & Johnson, 2007). his activation leads to the release of inflammatory cytokines, such as interleukin-6 (IL-6) (Welc *et al*. 2012) and interleukin-1β (IL-1β) (Lau, 2001). An overproduction of these cytokines can result in systemic inflammatory response syndrome (SIRS) (Balk, 2014), which has been linked to organ dysfunction, including liver and kidney damage (Garcia *et al*. 2022; Chapman *et al*. 2020; Geng *et al*. 2015; Wang *et al*. 2022). Given these findings from animal models, it is crucial to explore whether similar inflammatory responses occur in humans during extreme heat exposure, enabling the development of more effective mitigation strategies.

Current research on prolonged heat exposure primarily focuses on the susceptibility of older adults to SIRS, as indicated by elevated IL-6 levels in this population (Lee *et al*. 2024). his has led to the assumption that younger adults may tolerate prolonged heat exposure with minimal health consequences. However, this assumption is based on studies that present three critical limitations (Lee *et al*. 2024; Meade *et al*. 2023; McCormick *et al*. 2023). First, these studies largely involve dry heat exposure, which may not be directly applicable to humid heat conditions. It is well established that core temperature increases more rapidly in humid environments compared to dry heat (Cottle *et al*. 2022), suggesting that humid heat exposure could elicit a more pronounced inflammatory response and increase the risk of SIRS. Second, many of the studies involved compensable heat stress, with a minimal rise in *T_core_* (approximately 0.3 °C from baseline) at the end of the exposure. While this small increase was interpreted as insufficient to induce significant cytokine production, it is possible that such a minor temperature change is within the normal range of circadian fluctuation. Since these studies lasted 8–9 hours, the observed changes in *T_core_* may reflect normal circadian variations rather than a direct physiological response to heat stress. Therefore, the conclusion that younger adults are not at risk of SIRS under these conditions may be premature and warrants further investigation. Lastly, ad libitum fluid intake during these heat exposures may have mitigated the occurrence of SIRS, which could explain the lack of significant inflammatory responses observed in younger participants. However, these studies did not quantify participants’ hydration status before or after exposure, leaving the potential influence of hydration (euhydration vs. dehydration) on SIRS largely unexplored.

Given the direct threat that humid heatwaves pose to human health (Domeisen *et al*. 2023), it is critical to investigate how dehydration and euhydration influence the development of SIRS and organ function markers during prolonged extreme heat exposure. Fluid replacement is a widely accessible mitigation strategy that could significantly reduce the health impacts of extreme heat (Latzka *et al*. 1997; Goulet, 2013; Costello *et al*. 2018; Sawka *et al*. 2007). However, current studies primarily address the role of fluid replacement in compensable heat stress environments (Sawka & Montain, 2000), where heat stress is more easily managed by the body. The impact of fluid replacement on *T_core_* and SIRS in uncompensable heat stress conditions, where Tcore rises more significantly (Latzka *et al*. 1997), remains unclear. In these conditions, fluid replacement may not be sufficient to prevent SIRS, an issue that has not been extensively studied.

To address these gaps in the literature, this study aims to examine the effects of limited fluid intake versus fluid intake that fully replaces body weight loss on body temperature, SIRS, and organ function biomarkers (such as blood urea nitrogen [BUN] and alanine transaminase [ALT]) before and after 8 hours of extreme heat exposure in healthy young adults. We hypothesize that, at the end of the 8-hour exposure, younger adults will exhibit elevated levels of SIRS and organ function biomarkers (including BUN and ALT). Additionally, we hypothesize that fluid replacement, even when it fully replaces body weight loss, will not be sufficient to mitigate the occurrence of SIRS and organ dysfunction biomarkers following prolonged extreme heat exposure.

## Methods

### Participants

An a priori power analysis (G*Power version 3.1.9.4; Heinrich Heine University Düsseldorf, Düsseldorf, Germany) indicated that a minimum of five participants was required to address the main research question, using conventional α (0.05) and β (0.80) values, with an effect size of 0.859 (partial eta square), as reported by Mündel *et al*. 2010. The primary dependent variable for this calculation was IL-6 in response to heat stress compared to a temperate environment. Following approval from the institutional review board (XUST-IRB224002) and registration in the China Clinical Trial Database (ChiCTR2300071885), twelve non-acclimatized, healthy male participants (age: 25.0(1.6) years, body surface area (BSA): 1.9(0.1) m²) were recruited for this study. Although recruitment efforts targeted both males and females, only three female participants initially enrolled, but none completed all trials. Subsequent attempts to recruit female participants were unsuccessful.

### Experimental overview

All participants underwent two 8-hour extreme heat exposure trials under two hydration conditions: limited fluid intake (Dehydration, De) and fluid intake aimed at fully replacing fluid loss (Euhydration, Eu). Each trial began at 8:00 AM, with a five-day interval between trials to ensure adequate recovery. The order of trials was randomized and counterbalanced.

### Eight-hour extreme heat exposure

The heatwave simulation (40 ℃, 55% RH, heat index=58.6 ℃) (National Weather Service, 2023) selected for this study was based on conditions observed in several cities globally, such as the South and Midwest US during the summer, the 2017 Shanghai heatwave (heat index=56 ℃, July 2, 2017) (Rubin, 2019), the 2023 Mérida (Yucatán, Mexico) heatwave (heat index=58.3 ℃) (The Yucatan Times, 2023), the 2024 Rio de Janeiro heatwave (heat index=62.3 ℃) (Agence France Presse, 2024) and the 2020 Philippine heatwave (heat index=53-57 ℃, San Jose on April 20, 2020; Iba, Zambales, April 29, 2024) (Bella Cariaso, 2024). This simulation was chosen to represent uncompensable heat stress, as evidenced by continuous *T_core_* increases (Latzka *et al*. 1997) and aligns with projections of more frequent and humid heatwaves due to climate change (Mora *et al*. 2017).

To minimize seasonal acclimatization effects, all trials were conducted in early spring (late April to May), when the ambient temperature in Xi’an was approximately 30°C with 40% relative humidity, significantly lower than the experimental conditions. Before each heat exposure trial, participants voided their bladders to provide urine samples, ensuring their urine specific gravity (measured using a LH-Y12 meter; Lohand Biological, Hangzhou, China) was below 1.020 (Oppliger *et al*. 2005). After 15 minutes of seated rest, blood samples were collected via venipuncture, and participants weighed themselves nude. A rectal thermistor was self-inserted by participants, followed by attachment of other instruments. Participants then entered a climatic chamber (SEWTH-A-290H, Espec Environmental Instruments, Shanghai, China) to begin the 8-hour extreme heat exposure.

The exposure began with a 10-minute baseline period, followed by the 8-hour heat exposure. During the De condition, participants consumed 125 mL/hour of an electrolyte solution (lemonade with 0.4 mg zinc, 2.5 g carbohydrates, 25 mg sodium). In the Eu condition, participants consumed 375 mL/hour of the same solution. The 125 mL/hour intake was based on average adult consumption in China and the United States (Ma et al., 2012; Pokin et al., 2010). Participants engaged in normal office activities while seated throughout the trial. To standardize dietary intake, all participants were provided with a standard Big Mac burger (550 kcal). At the end of the 8-hour exposure, participants voided their bladders for a urine sample, weighed themselves nude, and had blood samples taken again via venipuncture. Body temperatures (rectal and skin) and heart rates were continuously recorded using data loggers, while blood pressure and thermal perceptions were measured every 60 minutes and 30 minutes, respectively.

### Measurements

#### Blood preparation

Venous blood samples were collected into vacutainers containing clot activator (Additive-free vacuum blood collection tubes, Kangwei Shi Medical Technology, Shijiazhuang, China). After clotting, samples were centrifuged at 4 ℃ and 3000g for 20 minutes. The resulting serum was aliquoted into Eppendorf tubes (2 mL centrifuge tubes, BKMAM, Changde, China) and stored at −80℃ for later analysis. Whole blood samples were also collected for plasma isolation using vacutainers containing EDTA (heparin tubes, Kangwei Shi Medical Technology, Shijiazhuang, China) as an anticoagulant. After centrifugation at 4°C and 3000g for 15 minutes, the plasma layer was carefully extracted, transferred into Eppendorf tubes (2 mL centrifuge tubes, BKMAM, Changde, China), and stored at −80°C for further analysis.

### Inflammatory cytokine markers

Inflammatory cytokines, including interlukin-4 (EK104-03), interlukin-6 (EK106HS-02), interlukin-1β (EK101BHS -01) and interferon-γ (EK180HS-AW1) were measured using enzyme-linked immunosorbent assay (ELISA) kits from Lianke Biotech Co. Ltd. (Hangzhou, China). The assays had the following intra-assay coefficients of variation (CVs): IL-4 (4.5%), IL-6 (4.7%), IL-1β (4.5%), and IFN-γ (4.3%). Sensitivities for these cytokines were 0.48 pg/mL, 0.02 pg/mL, 0.02 pg/mL, and 0.04 pg/mL, respectively.

### Oxidative stress, intestinal permeability and cortisol levels

Oxidative stress markers were measured as follows: malondialdehyde (MDA) using a kit from Nanjing Jiancheng Bioengineering Institute (2.3% intra-assay CV, sensitivity 0.5 nmol/mL), and superoxide dismutase (SOD) using a Servicebio kit (3% intra-assay CV, sensitivity 0.2 U/mL). Intestinal permeability was assessed by measuring intestinal fatty acid-binding protein (IFABP; Youersheng, Wuhan, China), and cortisol levels were quantified using an ELISA kit (Youersheng, Wuhan, China).

### Organ function markers

Blood urea nitrogen (BUN) and alanine transaminase (ALT) were measured to assess kidney and liver function, respectively. BUN was analyzed using an enzyme-labeled detector (Epoch, BioTeK, Winooski, USA), and ALT was measured using an automatic biochemistry analyzer (Chemray 800, Radu Life Sciences Shenzhen, Shenzhen, China).

### Electrolyte concentrations

Electrolyte concentrations (Na⁺, K⁺ and Cl⁻) were measured using an automated ion-selective electrode (ISE) analyzer (BKE-7, Biobase Group, Shandong, China). Calibration and quality controls were performed before each batch analysis, ensuring precision and accuracy. Results were reported in millimoles per liter (mmol/L). Three duplicate measurements were performed for validation, with intra- and inter-assay CVs maintained below 5%.

### Whole blood cell counts

Capillary blood samples were analyzed for leukocytes, hemoglobin, and hematocrit using an automatic hematology analyzer (Getein BHA-3000, Biotech Inc., Nanjing, China). All markers were corrected for plasma volume changes using the method described by Dill and Costill (1974).

### Cardiovascular measurements

Heart rate was continuously recorded using R-R detection (Polar Vantage XL, Polar Electro, Kempele, Finland). Blood pressure was measured using an automated monitor (YE660CR, Yuwell, Jiangsu, China). Measurements were taken in duplicate at rest, before and after meals, and hourly during the exposure. Mean arterial pressure (MAP) was calculated as diastolic blood pressure + 1/3 pulse pressure.

### Body temperatures

Core temperature *T*_core_ was measured using a rectal thermistor (YSI-401, USA; accuracy: 0.1 ℃), while mean skin temperature (*T*_sk_) was measured at four sites using iButtons (DS1922L, Vadisen Electronic Technology, Shanghai, China). The regional-weighted mean *T*_sk_ was calculated using the equation of Ramanathan (1964).

### Perceptions

Thermal comfort (7 points scale, −3: very uncomfortable to +3 very comfortable, Corgnati *et al*. 2007), thermal sensation (7 points scale, −3: very cold to +3 very hot, ASHRAE, 2020), wetness perception (7 points scale, −3: very wet to +3 very dry, Filingeri *et al*. 2015), thirst sensation (7 points scale, 1: not thirsty at all to 7: very very thirsty, Adams *et al*. 2020) and psychological stress (10 points scale, 0: no psychological stress to 10 very severe psychological stress, modified from Williams *et al*. 2010) were recorded every 30 minutes throughout the eight-hour extreme heat exposure.

### Statistical analysis

All statistical analyses were performed using SPSS software for Windows (IBM SPSS Statistics 20, NY, USA), and graphical representations were generated using GraphPad Prism (Prism version 7.00, GraphPad Software). The homogeneity of variance was assessed using Levene’s test, and data normality was evaluated using the Kolmogorov-Smirnov test. To assess the effect of fluid intake on inflammatory and thermoregulatory responses at different time points, a two-way repeated measures ANOVA was performed, considering the factors of hydration condition (Dehydration vs. Euhydration) and time point. When significant main or interaction effects were identified, post hoc pairwise comparisons were conducted using paired t-tests with Bonferroni adjustments, where appropriate. All data are presented as means ± standard deviation (SD), unless otherwise specified. Statistical significance was defined as a *P*-value of < 0.05.

## Results

### Quantification of hydration level

Urine specific gravity (USG) was significantly higher in the Dehydration (De) condition compared to the Euhydration (Eu) condition at the end of the exposure (1.030 ± 0.004 vs. 1.020 ± 0.020, respectively, *P*<0.01). Additionally, the dehydration rate was significantly greater in the De condition (3.0 ± 0.4%) compared to the Eu condition (no dehydration) (*P*<0.01). These results confirm that dehydration was more pronounced in the De condition than in the Eu condition (Table 1).

**Table 1.** Physiological, biochemical, and perceptual responses, and plasma electrolyte concentrations before (Pre) and after (Post) the eight-hour daylong heat exposure in the Dehydration (De) and Euhydration (Eu) trials. *Indicates a significant difference compared to pre-exposure; #Indicates a significant difference between Eu and De conditions.

|  | De (dehydration) |  | Eu (euhydration) |  |
| --- | --- | --- | --- | --- |
| **Cardiovascular** | **Baseline** | **End** | **Baseline** | **End** |
| Systolic blood pressure (mmHg) | 119<br>(7) | 114<br>(8)* | 121<br>(9) | 114<br>(7)* |
| Diastolic blood pressure (mmHg) | 72<br>(7) | 62<br>(8)* | 71<br>(5) | 64<br>(6)* |
| Mean arterial pressure (mmHg) | 87<br>(7) | 79<br>(7)* | 87<br>(6) | 80<br>(6)* |
| Heart rate (beats/min) | 87<br>(9) | 118<br>(9)* | 90<br>(8) | 117<br>(10)* |
| **Biochemistry markers** | **Baseline** | **End** | **Baseline** | **End** |
| Haematocrit (%) | 46<br>(1.9) | 47<br>(2.1)* | 46.4<br>(3.05) | 45.7<br>(2.5)* |
| Urine specific gravity (g/mL) | 1.013<br>(0.006) | 1.03<br>(0.004)* | 1.016<br>(0.005) | 1.02<br>(0.02)*# |
| Leukocytes ($10^9/L$) | 6.7<br>(1.2) | 12.9<br>(2.2)* | 6.1<br>(0.9) | 13.1<br>(1.9)* |
| Neutrophils ($10^9/L$) | 3.5<br>(0.7) | 9.1<br>(2.1)* | 3.3<br>(0.5) | 9.0<br>(1.8)* |
| **Perceptions** | **Baseline** | **End** | **Baseline** | **End** |
| Thermal sensation | 1.8<br>(0.6) | 2.4<br>(0.4)* | 1.7<br>(0.4) | 2.2<br>(0.4)* |
| Thermal comfort | -0.7 | -2.4 | -1.0 | -2.0 |
|  | (0.5) | (0.4)<sup>*</sup> | (0.4) | (0.7)<sup>*</sup> |
| Wetness perception | -0.9 | -2.4 | -1.1 | -2.1 |
|  | (0.7) | (0.6)<sup>*</sup> | (0.4) | (0.5)<sup>*</sup> |
| Thirst sensation | 4.2 | 4.8 | 4.0 | 4.1 |
|  | (0.3) | (0.7)<sup>*</sup> | (0.0) | (0.2)<sup>*</sup> |
| Psychological stress | 0.3 | 6.0 | 0.3 | 5.0 |
|  | (0.5) | (1.0)<sup>*</sup> | (0.5) | (1.0)<sup>*</sup> |
| **Plasma electrolytes** | **Baseline** | **End** | **Baseline** | **End** |
| K<sup>+</sup> | 3.6 | 3.8 | 3.9 | 3.6 |
| (mmol/L) | (0.2) | (0.2)<sup>*</sup> | (0.1) | (0.1)<sup>*</sup> |
| Na<sup>+</sup> | 144.7 | 146.3 | 147.9 | 140.2 |
| (mmol/L) | (1.7) | (1.4)<sup>*</sup> | (1.7) | (1.1)<sup>*</sup> |
| Cl<sup>-</sup> | 102.9 | 105.3 | 102.6 | 101.4 |
| (mmol/L) | (1.0) | (0.8)<sup>*</sup> | (1.7) | (1.1) |
| **Dehydration** | **Baseline** | **End** | **Baseline** | **End** |
| Dehydration rate | - | 3.01 | - | Euhydrated |
| (%) |  | (0.5)<sup>*</sup> |  |  |
| Average sweating rate | - | 250.7 | - | 283.1 |
| (g/h) |  | (44.4)<sup>*</sup> |  | (89.8)<sup>*</sup> |

Note: -, not applicable.

### Body temperatures

Core temperature (*T*_core_) did not significantly differ between the De and Eu conditions (*P*=0.31). However, a significant interaction effect was observed between hydration condition and time points (Fig.1A, *P*<0.05). Specifically, *T*_core_ was higher in the De condition than in the Eu condition at the end of the exposure. Similarly, mean skin temperature (*T_sk_*) did not show a significant main effect between conditions, but an interaction effect was found between hydration condition and time points (Fig.1B, *P*=0.01). Post hoc comparisons revealed no significant differences in *T_sk_* between the De and Eu conditions at any time point.

**Figure 1.**
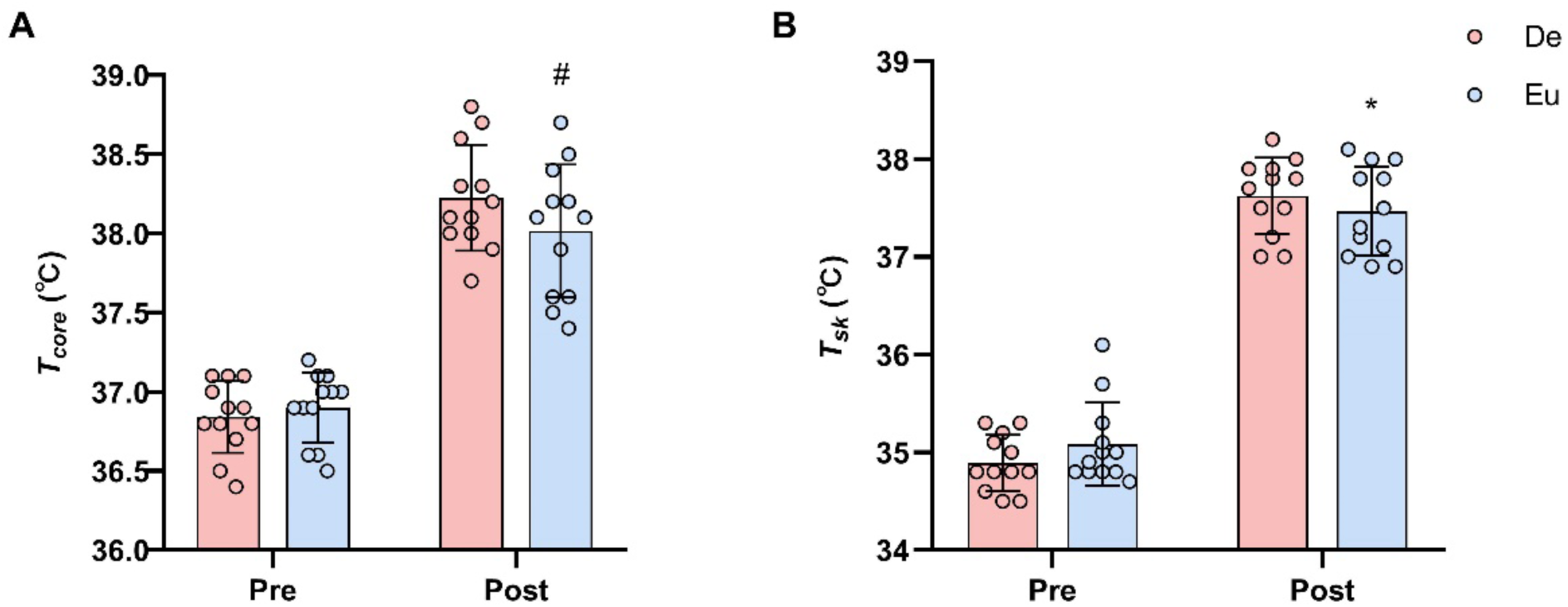
Core temperature (*T_core_*, Figure 1A) and mean skin temperature (*T_sk_*, Figure 1B) before (Pre) and after (Post) eight-hour extreme heat exposure in the Dehydration (De) and Euhydration (Eu) trials. *Indicates a significant difference compared to pre-exposure; #Indicates a significant difference between Eu and De conditions.

### Oxidative stress, gastrointestinal permeability, and cortisol levels

Both malondialdehyde (MDA) and superoxide dismutase (SOD) levels were significantly higher at the end of the exposure compared to pre-exposure levels in both the Euhydration (Eu) and Dehydration (De) conditions (Figs. 2A & 2B, all *P*<0.05). However, only SOD levels differed significantly between conditions (*P*<0.01), with a notable interaction effect between conditions and time points (*P*<0.01). Specifically, SOD levels were higher in the Eu condition than in the De condition at the end of the exposure.

**Figure 2.**
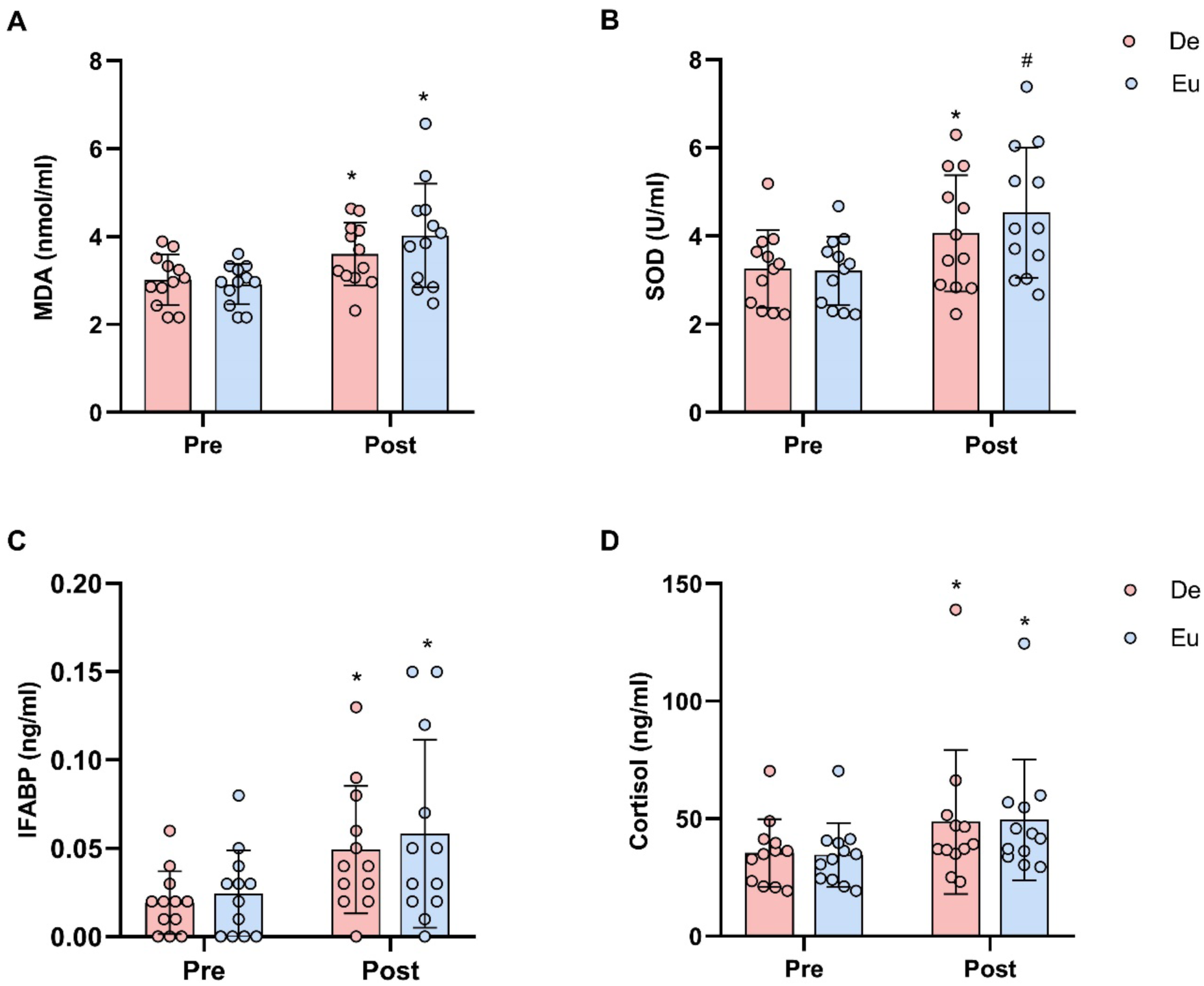
Malondialdehyde (MDA, Figure 2A), Super Oxide Dismutase (SOD, Figure 2B), gastrointestinal permeability (IFABP, Figure 2C), and cortisol (Figure 2D) before (Pre) and after (Post) the eight-hour heat exposure in the Dehydration (De) and Euhydration (Eu) trials. *Indicates a significant difference compared to pre-exposure; #Indicates a significant difference between Eu and De conditions.

Intestinal fatty acid-binding protein (IFABP) levels were also higher at the end of the exposure compared to pre-exposure levels in both conditions (Fig. 2C, *P*=0.013). However, IFABP levels did not differ significantly between conditions and showed no interaction effect between conditions and time points (*P*=0.37). Cortisol levels were significantly higher at the end of the exposure compared to pre-exposure levels in both the Eu and De conditions (*P*<0.01). However, cortisol did not differ between conditions (P = 0.98) and showed no interaction between conditions and time points (*P*=0.54) (see Fig. 2D).

### Systemic inflammatory response and visceral organ function markers

Leukocyte levels did not differ significantly between conditions (Table 1, *P*=0.51), although a significant interaction effect between conditions and time points was observed (*P*=0.048). Post hoc analysis revealed that leukocyte levels were significantly higher at the end of the exposure compared to pre-exposure in both conditions. In contrast, neutrophil levels were higher at the end of the exposure compared to pre-exposure, but there was no significant difference between conditions (*P*=0.53), and no interaction effect between conditions and time points (*P*=0.91).

Interleukin-6 (IL-6) levels differed significantly between conditions (Fig. 3A, *P*<0.01) and showed a significant interaction effect between conditions and time points (*P*<0.01). Specifically, IL-6 levels were higher at the end of the exposure compared to pre-exposure, and levels in the Eu condition were higher than those in the De condition at the end of the exposure. Other inflammatory markers, such as IL-1β and IFN-γ, were also elevated at the end of the exposure compared to pre-exposure (all *P*<0.01), but no significant differences were observed between conditions (all *P*>0.40). Furthermore, these markers showed no interaction effects between conditions and time points (all *P*>0.40).

**Figure 3.**
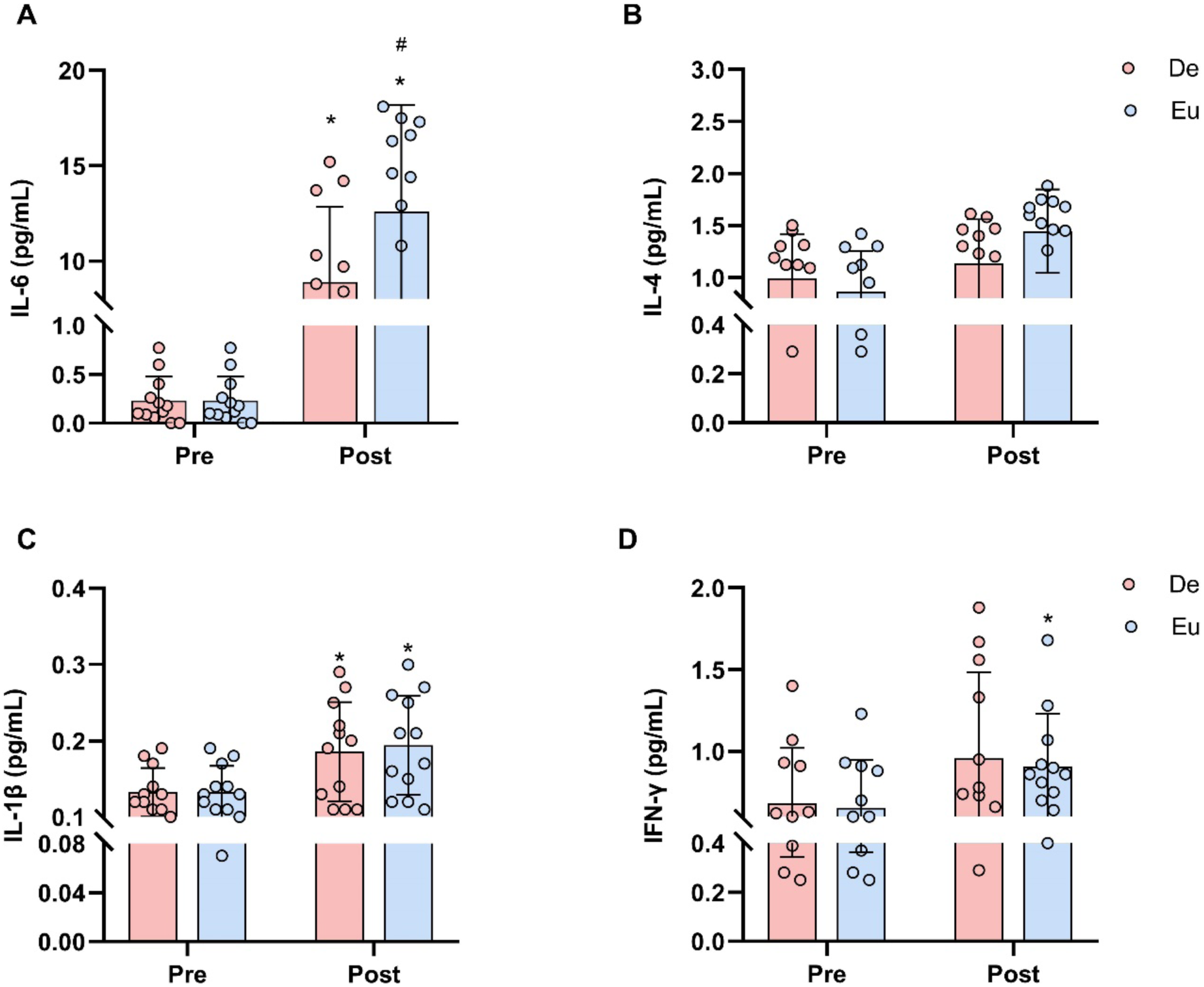
Pro-inflammatory cytokine responses: interleukin-6 (Figure 3A, interleukin-4 (Figure 3B), interleukin-1β (Figure 3C), and interferon-γ (Figure 3D) before (Pre) and after (Post) the eight-hour daylong heat exposure in the Dehydration (De) and Euhydration (Eu) trials. *Indicates a significant difference compared to pre-exposure; #Indicates a significant difference between Eu and De conditions.

Both alanine aminotransferase (ALT) and blood urea nitrogen (BUN) levels were significantly higher at the end of the exposure compared to pre-exposure levels (Fig. 4, all *P*<0.01). However, only ALT levels showed a significant difference between conditions (*P*=0.03), with a significant interaction effect between conditions and time points (*P*=0.03). Specifically, ALT levels were higher in the Eu condition than in the De condition at the end of the exposure.

**Figure 4.**
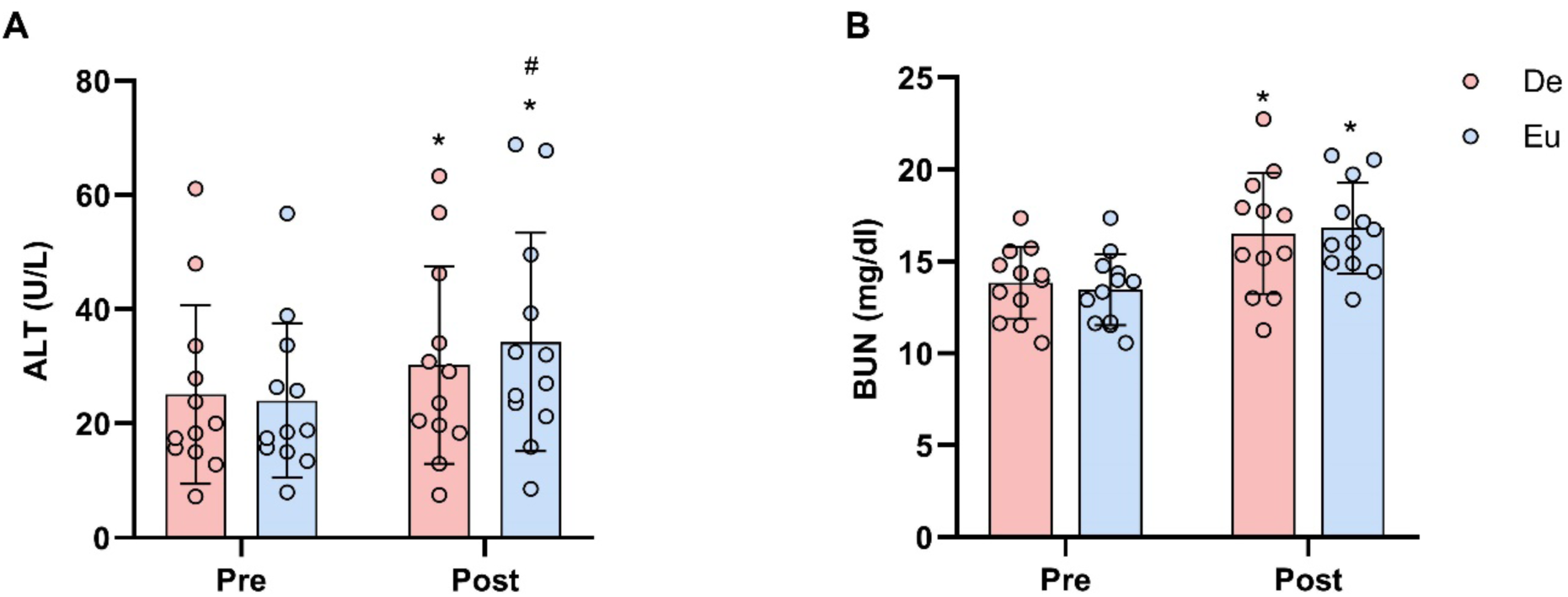
Markers of visceral organ function: liver alanine aminotransferase (ALT, Figure 4A] and kidney blood urea nitrogen (BUN, Figure 4B) before (Pre) and after (Post) the eight-hour daylong heat exposure in the Dehydration (De) and Euhydration (Eu) trials. *Indicates a significant difference compared to pre-exposure; #Indicates a significant difference between Eu and De conditions.

### Cardiovascular responses

Systolic blood pressure, diastolic blood pressure, and mean arterial pressure were all lower at the end of the exposure compared to pre-exposure levels in both the Euhydration (Eu) and Dehydration (De) conditions (all *P*<0.01). However, these measures did not differ significantly between the two conditions (all *P*>0.70) and showed no interaction effects between conditions and time points (all *P*>0.10). In contrast, heart rate was significantly higher at the end of the exposure compared to pre-exposure (*P*<0.01), but there were no significant differences between conditions (*P*=0.92), and no interaction effect was observed between conditions and time points (Table 1, *P*=0.35).

### Plasma electrolyte concentrations

In the De condition, plasma concentrations of potassium, sodium, and chloride increased by 0.2 mmol/L, 1.6 mmol/L, and 2.4 mmol/L, respectively (all *P*<0.05). In contrast, in the Eu condition, plasma concentrations of potassium and sodium decreased by 0.3 mmol/L and 7.7 mmol/L, respectively (all *P*<0.05, see Table 1).

### Perceptions

Thermal comfort differed significantly between conditions and showed a significant interaction effect between conditions and time points (Table 1, *P*=0.014). Specifically, participants reported feeling more comfortable in the Eu condition than in the De condition post-exposure. Thirst sensation also differed significantly between conditions (*P*<0.05) and time points (Table 1, *P*=0.02). Participants felt less thirsty in the Eu condition than in the De condition at the end of the exposure. In contrast, thermal sensation did not differ significantly between conditions (*P*=0.44) and showed no interaction effect between conditions and time points (*P*=0.42). Psychological stress was not significantly different across conditions (*P*=0.20) and revealed no interactional effect between conditions and timepoints (*P*=0.18).

## Discussion

This study yielded two major findings: (i) Systemic inflammatory response syndrome (SIRS) was evident at the end of the 8-hour heat exposure, resulting in an increased risk of acute kidney and liver dysfunction; (ii) Fluid replenishment during this exposure exacerbated SIRS and escalated the risk of liver dysfunction, despite a lower *T_core_*. These findings support our hypothesis and indicate that replenishing fluids to match total body weight loss during prolonged extreme heat exposure may worsen SIRS and increase the risk of liver dysfunction in healthy Asian men.

The presence of SIRS at the end of the 8-hour extreme heat exposure was evidenced by elevated levels of IL-6, IL-1β, IFN-γ, leukocytes, and neutrophils in both the Eu and De conditions (Figure 3, Table 1). These findings contrast with previous studies (Lee *et al*. 2024; Meade *et al*. 2023; McCormick *et al*. 2023), which did not observe SIRS following 8-9 hours of heat exposure. The discrepancy could be attributed to the greater rise in *T_core_* in our study (1.5 ℃ vs. 0.3 ℃, Figure 1), which may have led to increased gastrointestinal leakage of IFABP (March *et al*. 2017; Ogden *et al*. 2020; Leon & Helwig, 2010; Lambert, 2004; Pires *et al*. 2003; Lang *et al*. 2003) and greater cellular stress on toll-like receptor 4 (TLR4) in tissues (Meador *et al*. 2008). This cellular stress likely upregulated heat shock factor 1 (HSF1), resulting in excessive IL-6 production (McLoughlin *et al*. 2003; Leon & Helwig, 2010). Our data partially support this hypothesis, as we observed elevated IFABP levels at the end of the exposure, along with higher oxidative stress (Figure 2). These findings indirectly confirm heat-induced stress on TLR4, though the absence of direct MAPK and TLR4 pathway analysis necessitates further research to confirm this mechanism. Another possible contributor to the observed SIRS is psychological stress. Extreme heat exposure may trigger a rise in sympathetic nerve activity, promoting the migration of monocytes and macrophages into systemic circulation and subsequently increasing IL-6 and IL-1β secretion (Yamakawa *et al*. 2009; Maes *et al*. 1998). This is supported by our data, which show that cortisol concentrations and psychological stress were significantly higher following the 8-hour heat exposure in both Eu and De trials (Figure 2, Table 1). Our findings are consistent with animal studies that found IL-6 concentrations to be elevated compared to other pro-inflammatory cytokines after extreme heat exposure (Welc *et al*. 2013). Elevated IL-6 likely triggers the secretion of IL-1β and IFN-γ (Leon & Helwig, 2010), ultimately contributing to organ dysfunction and damage (Chapman *et al*., 2021; Giercksky *et al*., 1999). This mechanism is supported by our data, which show higher IL-6 concentrations compared to IL-1β and IFN-γ, accompanied by increased BUN and ALT levels (Figures 3 & 4).

Interestingly, fluid intake during the 8-hour extreme heat exposure did not mitigate the acute SIRS response. On the contrary, it exacerbated SIRS and increased the risk of liver dysfunction, as evidenced by higher IL-6 and ALT levels in the Eu condition compared to De, despite a lower *T_core_* (Figures 3 & 4). These findings suggest that pro-inflammatory cytokine secretion is not solely dependent on *T_core_*. We propose that excessive fluid intake may have induced hypotonic osmotic stress, which occurs when the extracellular fluid has a lower solute concentration than the intracellular fluid. This osmotic imbalance leads to fluid influx into cells, causing cell swelling, increased cellular stress, and, potentially, cell lysis or apoptosis (Groulx *et al*. 2006, Jäckle *et al*. 2001). In our study, hypotonic stress was evident due to excessive fluid intake diluting extracellular solute concentrations compared to intracellular fluid (see Table 1 for plasma electrolytes). Consequently, intracellular fluid accumulation likely caused swelling of liver and kidney cells, contributing to the observed organ dysfunction. This cellular stress could have triggered oxidative stress, characterized by increased production of reactive oxygen species (ROS), which can exacerbate tissue damage (Hu *et al*. 2017,). The higher SOD levels observed in the Eu condition support this hypothesis, as the body attempts to counteract oxidative damage. Furthermore, hypotonic osmotic stress may activate endocytic activity in macrophages via ClC-3 channels (Yan et al., 2014), potentially leading to greater IL-6 secretion (Xiang et al., 2016). This mechanism could explain the elevated IL-6 levels in the Eu condition compared to De (Figure 3). Additionally, hypotonic osmotic stress might exacerbate liver injury, as acute or chronic stress can increase hepatocyte damage, leading to elevated ALT concentrations (Ji *et al*., 2024). The higher ALT levels in the Eu condition may therefore reflect direct liver cell damage due to cellular swelling (Figure 4). Taken together, these findings suggest that excessive fluid intake induced hypotonic osmotic stress, worsening cellular damage in the liver and kidneys, despite a lower *T_core_*, and thereby increasing the risk of organ dysfunction. This underscores the delicate balance required in hydration strategies during prolonged heat exposure, as both dehydration and overhydration pose significant health risks.

### Limitations

While this study effectively highlights the effects of fluid intake on SIRS during 8 hours of extreme heat exposure, there are two key limitations. First, we did not include female participants, leaving the effect of sex differences on SIRS during extreme heat exposure unresolved. As Bouchama *et al*. (2023) utilized whole-genome transcriptomics to identify sex-specific molecular signatures of heat stroke, similar studies should be conducted to explore the differential response to heat in males and females. Second, we did not investigate the ERK1/2 and JNK MAPK pathways, particularly the role of c-Fos and c-Jun in the early stress-induced cytokine response. Future research should explore these pathways to better understand the cellular mechanisms driving the inflammatory response to heat exposure.

### Practical significance

Based on our findings, we propose several practical recommendations regarding hydration strategies during prolonged extreme heat exposure. First, hydration should be personalized based on sweat rate monitoring. Instead of a fixed fluid replacement approach, individuals should measure their sweat rate by tracking body weight changes before and after exposure. A loss of 1–2% body weight indicates dehydration, and fluid replacement should match hourly sweat loss, typically ranging from 500 to 1500 mL/hour. Second, electrolyte supplementation should be incorporated into hydration strategies. Excessive fluid intake without adequate electrolyte replacement can cause hyponatremia and exacerbate hypotonic stress. Fluids containing 500–700 mg/L of sodium, with 1– 2 grams of sodium chloride per liter, should be used to maintain electrolyte balance and prevent cell damage. Third, monitoring for signs of overhydration is essential, as excessive fluid intake can exacerbate SIRS and liver injury. Occupational workers or athletes should limit fluid intake to no more than 1.2 times their hourly sweat loss, and early signs of overhydration, such as bloating, nausea, or dizziness, should be closely monitored.

Additionally, osmolarity monitoring using urine specific gravity (USG) can assist in tracking hydration status in real time. A USG value between 1.010 and 1.020 indicates proper hydration, while values below this suggest overhydration. Regular body mass checks before and after shifts can help maintain hydration within 1-2% of baseline body weight. Fluid intake should also be spaced out, with small, steady sips (125-250 mL every 15-20 minutes), rather than large intakes at once, to prevent osmotic stress. Post-exposure recovery should focus on replenishing both fluids and electrolytes lost during heat exposure, using options like sports drinks or salty snacks. Monitoring psychological and physiological responses, including thirst, thermal comfort, and stress, can provide real-time cues for adjusting hydration levels as needed. Finally, fluid management should remain dynamic, with adjustments based on environmental conditions and individual needs to optimize health outcomes during heat exposure.

## Conclusions

This study provides compelling evidence that fluid replacement during eight-hour extreme heat exposure does not mitigate SIRS and may in fact exacerbate liver dysfunction. Despite the expected benefits of fluid intake in maintaining core temperature, excessive hydration leads to hypotonic osmotic stress, triggering a pro-inflammatory response characterized by elevated IL-6 and oxidative stress markers. These findings emphasize the need for more nuanced hydration strategies that balance fluid and electrolyte intake to avoid both dehydration and overhydration. Future research should explore sex differences and investigate molecular pathways such as the MAPK signaling cascade to further elucidate the mechanisms driving heat-induced SIRS and associated organ dysfunction.

## Additional information

### Data availability statement

The data supporting the present findings are available from the corresponding author upon reasonable request. Due to privacy or ethical restrictions, the data is not publicly available.

### Author contributions

F.W. and T.H.L. conceived and designed experiments. All authors performed experiments. F.W. and T.H.L. analysed the data. F.W. and T.H.L. drafted the manuscript. All authors were involved in revising the manuscript. All authors have read and approved the final version of this manuscript and agree to be accountable for all aspects of the work in ensuring that questions related to the accuracy or integrity of any part of the work are appropriately investigated and resolved. All persons designated as authors qualify for authorship, and all those who qualify for authorship are listed.

